# Side effects and the need for secrecy: characterising discontinuation of modern contraception and its causes in Ethiopia using mixed methods

**DOI:** 10.1101/140004

**Authors:** Alexandra Alvergne, Rose Stevens, Eshetu Gurmu

## Abstract

**Background:** Contraceptive discontinuation is a major barrier to reducing global unmet needs for family planning, but the reasons why women discontinue contraception are poorly understood. Here we use data from Ethiopia to investigate (i) the magnitude of contraceptive discontinuation in 2005-2011, (ii) how the risk of discontinuation varies with method type and education level and (iii) the barriers to continuation. Our main hypothesis is that contraceptive discontinuation is driven by the experience of physiological side-effects associated with the use of hormonal contraception, rather than a lack of education.

**Methods:** We used a mixed methods explanatory sequential design to explain the quantitative results in more details through the qualitative data. First, we analysed quantitative data from the 2011 Ethiopian Demographic and Health Survey to study patterns of contraceptive discontinuation and method choice using multilevel multiprocess models. Second, we conducted semi-structured interviews and focus group discussions in the 3 most populated regions of Ethiopia with individuals of reproductive age and health professionals.

**Results:** The analysis of EDHS data shows that the rate of discontinuation has not reduced in the period 2005-2011 and remains high. Discontinuation mainly takes the form of abandonment, and is a function of method type and wealth but not of educational level. Interviews with women and health professionals reveal that the experience of debilitating physiological side effects, the need for secrecy and poverty are important barriers to continuation.

**Conclusions:** Our findings together suggest that physiological and social side-effects of contraceptive use, not education, are the root causes of contraceptive discontinuation in Ethiopia. We argue that to tackle discontinuation due to side-effects, dispelling misconceptions through educating women is not addressing the root causes of discontinuation, and that priority should be given to both engaging men and questioning the appropriateness of medical technology to the physiology of Ethiopian women, especially those living in poverty.

## Background

### Contraceptive discontinuation as a major determinant of unmet needs

Unwanted fertility leads to increased rates of abortion and maternal mortality, and acts as a major barrier to improving individual health, gender equity, family wellbeing and national development [1–4]. Thus, tackling unmet need for contraception, i.e. the proportion of women wishing to limit or postpone child birth but who are not using contraception [5], has recently been highlighted as a key global health issue [6]. In low-resource countries, 1 in 4 women has an unmet need for family planning and 79% of unintended pregnancies occur among these women [2]. Yet, a main obstacle to reduce unwanted fertility is the discontinuation of modern contraception [7–9], a currently underappreciated proportion of unmet need. On average, over a third of women who start using a modern contraceptive method stop using it within the first year, and over a half stop before two years [9]. Following Jain’s claim that programs would do better to concentrate on retaining existing users rather than focus on recruiting new clients [10, 11], recent reports have suggested that tackling contraceptive discontinuation is central to achieving better demographic outcomes [7, 8]. To date, however, there is little valid literature on either the reasons for discontinuation [12] or the programmatic interventions explicitly designed to reduce discontinuation [9].

### The reasons for contraceptive discontinuation

Why women discontinue contraception is multi-faceted, but several reviews of DHS data have demonstrated that side effects and health concerns associated with the use of hormonal contraceptives are major reasons for discontinuation [7–9, 11, 13]. In a review of oral contraceptive (OC) discontinuation in 19 developing countries, Ali & Cleland [7] found that the dominant reason for terminating OC use within 12 months is dissatisfaction with the method, predominantly side-effects and health concerns. These account for a median of 28% of all reasons for discontinuation, reaching as high as 40% in Bolivia [13]. Yet, most current studies are based on DHS data, which only records one general reason for discontinuation and one for non-use. Answers are then often grouped into large ambiguous categories such as ‘health concerns’ or ‘opposition to use’ making it difficult to understand the nuances behind the reasons [14]. There have been several qualitative studies on attitudes towards contraceptive use that cite health concerns as a barrier to contraceptive use and continuation [15–18], but, similarly to DHS data, the distinction between unsubstantiated fears and the experience of side effects is not systematically articulated. This is problematic because it leads to the assumption that concerns about side-effects are grounded in myths and misconceptions that can be alleviated by counselling and education [5, 8, 9], disregarding the possibility that side-effects may be intolerable [19].

### Hormonal contraceptives and physiological side-effects

The experience of physiological side-effects associated with the use of hormonal contraception has been hypothesized to result from a mismatch between a woman’s endogenous hormonal levels and the dosage of contraceptives [20]. Hormonal contraceptives mainly work by inhibiting ovulation through mimicking the post-ovulatory phase of the menstrual cycle and the early stage of pregnancy, that is, by releasing synthetic ovarian steroids (progestogen and/or estrogen) so that to initiate a negative feedback that prevent the surge of the hormones responsible for ovulation [21]. The dosage must be high enough so that to prevent ovulation, but not too high so as not to lead to side-effects (nausea, vomiting, headache, etc.). This balance is likely to be offset in the case of many women living in the developing world because the contraceptives used by family planning programs are generally trialled on Western, industrialized and non-seasonal populations, and thus contain high levels of ovarian steroids. Indeed, there is widespread evidence that female reproductive functioning is highly variable between populations of different subsistence modes [20, 22, 23], and within populations, between seasons, socio-economic classes and migration status [24, 25]. This suggests that health concerns might be rooted in the experience of side-effects, independently of education level.

### Contraceptive discontinuation in Ethiopia

In Ethiopia, the focus of this paper, contraceptive prevalence has increased nine-fold from 1990 to 2011 and to date, it continues to rise. The increase in the number of contraceptive users speaks, to some extent, to the success of the Health Extension Program launched by the Federal Ministry of Health (FMOH) of Ethiopia in 2003 to reach the under-served in rural areas. This community-based service provision based on the diffusion model [26] has led to the training of more than 30,000 health extension workers [27, 28]. Yet, some family planning outcomes fall far from the targets set by the international partnership Family Planning 2020. For instance, while the Ethiopian government committed to reducing total fertility rate (TFR) to 4 and increasing contraceptive prevalence to 69% by 2015, by 2016, TFR is at 4.6 (5.2 and 2.3 in rural and urban areas, respectively) and contraceptive prevalence is at 35.9%. Further, based on the analysis of data from Demographic and Health Surveys (DHS) from 25 countries including DHS data collected in Ethiopia in 2005, it was found that both the level of unmet need for contraception (33.8%) and contraceptive discontinuation due to method-related dissatisfaction were the highest in Ethiopia [8]. Although the picture has somewhat improved, with unmet needs down to 25.3% among married women in 2011 and 22.3% in 2016, there is scope for further decreasing unmet needs for contraception. A promising avenue might be to make the tackling of contraceptive discontinuation a priority.

### Aim of the study

This paper seeks to characterize and improve understanding of the determinants of contraceptive discontinuation in Ethiopia, using both quantitative and qualitative data. Specifically, we aimed to investigate (1) the magnitude of contraceptive discontinuation at the country level in 2011, and how it compares to the situation observed in 2005, (ii) the roles of method type and education level in modulating the risk of contraceptive discontinuation and (iii) the barriers to contraceptive continuation. Our main hypothesis is that contraceptive discontinuation is driven by the experience of physiological side-effects associated with the use of hormonal contraception, rather than a lack of education.

## Methods

### Study Design and Population

First, we conducted a statistical analysis of a nationally representative sample of women of reproductive age using the Ethiopia 2011 DHS Survey, focusing on unsterilized women who have ever used modern contraception in the preceding 5 years of the survey. Because method choice and discontinuation are not independent, i.e. women who might want to discontinue are more likely to choose short over long acting methods [11], we used a multilevel and multiprocess model [29] to avoid overestimating the discontinuation of short-acting methods. Second, given the Ethiopian 2011 DHS data does not inform on the reasons for discontinuation, a qualitative study on the experience of contraceptive use was run to supplement the quantitative analysis. Interviews and focus group discussions were conducted in 2013 in the three most populated regions in the country (the rural Arsi Administrative Zone of Oromia Region, the Sidama Administrative Zone of SNNP Region and the North Shewa Administrative Zone of Amhara Region). These regions were chosen to provide a variety of different ethnicities interviewed with different perspectives on contraceptive use.

### Analysis of the Ethiopian 2011 Demographic and Health Survey

The ethical procedure associated with DHS data collection can be found on the DHS website [30]. The contraceptive histories are collected using a “calendar” that records monthly contraceptive status during the 5 years preceding the survey. The 2011 EDHS dataset includes 16,615 women, married and unmarried, aged 15-49, among which 30.56% have ever used a contraceptive method (29.15% have ever used a modern contraceptive method and 17.93% are currently using a method). Following previous analyses [29], we distinguished 3 types of discontinuation: failure (unintentional) as opposed to abandonment and switch (intentional). Failure corresponds to a situation in which a woman reports becoming pregnant while using a given method (so it includes failure of the method itself and failure to use it). Abandonment corresponds to the absence of use for more than a month and may lead to an immediate risk of unwanted pregnancy. Switch corresponds to a switch from one method to another within a month and as defined here, does not lead to periods during which a woman is unprotected. We focused on modern methods and distinguished 4 different types (1) oral contraceptives (OC), (2) injectables, (3) IUD and implants (long term methods which require health workers to remove it) and (4) condoms. Sterilized women (1.4% in 2011) were excluded, as this contraceptive method is non-reversible and when discussing condoms, we are referring to male as opposed to female condoms.

#### 1. Modelling contraceptive discontinuation and method choice: an overview

To analyse the risk of contraceptive discontinuation over time, the data have been converted into a period-person dataset, considering the unit of analysis to be an episode of contraceptive use. An episode is defined as a continuous period of use of a specific method. Once an individual has experienced one type of discontinuation, the individual is removed from the observations; the survival time for the competing risks are latent and we only observe *T* = min (*T*(failure), *T*(switch), *T*(abandon)) or the censoring time if no discontinuation has occurred [29]. In this analysis we concentrate on women who have ever-used a modern contraceptive method and who have not undergone sterilization as this method cannot be discontinued sterilized (*N* = 4703). The analysis includes data on 7022 episodes of use contributed by 4703 ever-users. Within those episodes of contraceptive use, 3965 events of discontinuation were observed (56.5%). In the case of discontinuation events, 64.6% correspond to abandonment; 15.9% correspond to failure and 19.5% correspond to switching.

Following Steele & Curtis [29], the analysis proceeds in 2 steps using the aML statistical software (http://www.applied-ml.com). First, as there are 3 types of discontinuation, we used a competing risk hazard model, with one equation per risk of discontinuation. The model allows for unobserved heterogeneity between women (i.e. random effects) that may influence the duration of each episode. To investigate if the risks are correlated (e.g. the risk of failure is not independent from the risk of abandonment if a woman wants to avoid pregnancy), the random effects must be correlated across the different types of discontinuation; and thus the 3 equations must be estimated simultaneously. Second, because some women who are more likely to discontinue are more likely to use short-acting methods, the analysis must consider that method choice is a potential endogenous variable. To address this issue, we use a multilevel multiprocess model that simultaneously models the processes of contraceptive choice and contraceptive discontinuation. In all analyses, our variables of interest were method type and a woman’s educational level. We all also included covariates that have been previously found to relate to contraceptive discontinuation, i.e. household socio-economic status, area of residence, religion and the age of the women at the start of the contraceptive episode. We did not include marital status because it has likely changed over the 5 years’ calendar period (in the data 87% of women are living with partner in 2011). Each step of the analysis is further detailed below.

#### 2. Standard multilevel hazards models for competing risks

First, we modelled discontinuation using a competing risk hazard model. We built one model per type of discontinuation step by step: (i) estimation of a simple Gompertz hazard model, without covariates; (ii) adding heterogeneity by incorporating residuals following a univariate normal distribution; (estimating the unobserved woman-level effect). The inclusion of random effects enables to account for unobserved woman-level characteristics that influence the hazard of discontinuation at each month of a given episode and for each episode. The 3 types of discontinuation were modelled jointly (3 equations) to allow for the possibility that random effects are correlated across equations. Then, we ran the 3 models simultaneously, first with uncorrelated random effects (e.g. assuming that the risks do not compete) and then with correlated random effects (assuming that there are competing risks). We found that those effects were not correlated (Table 2) thus single hazard models (one per type of discontinuation) were subsequently modelled.

**Table 1:**
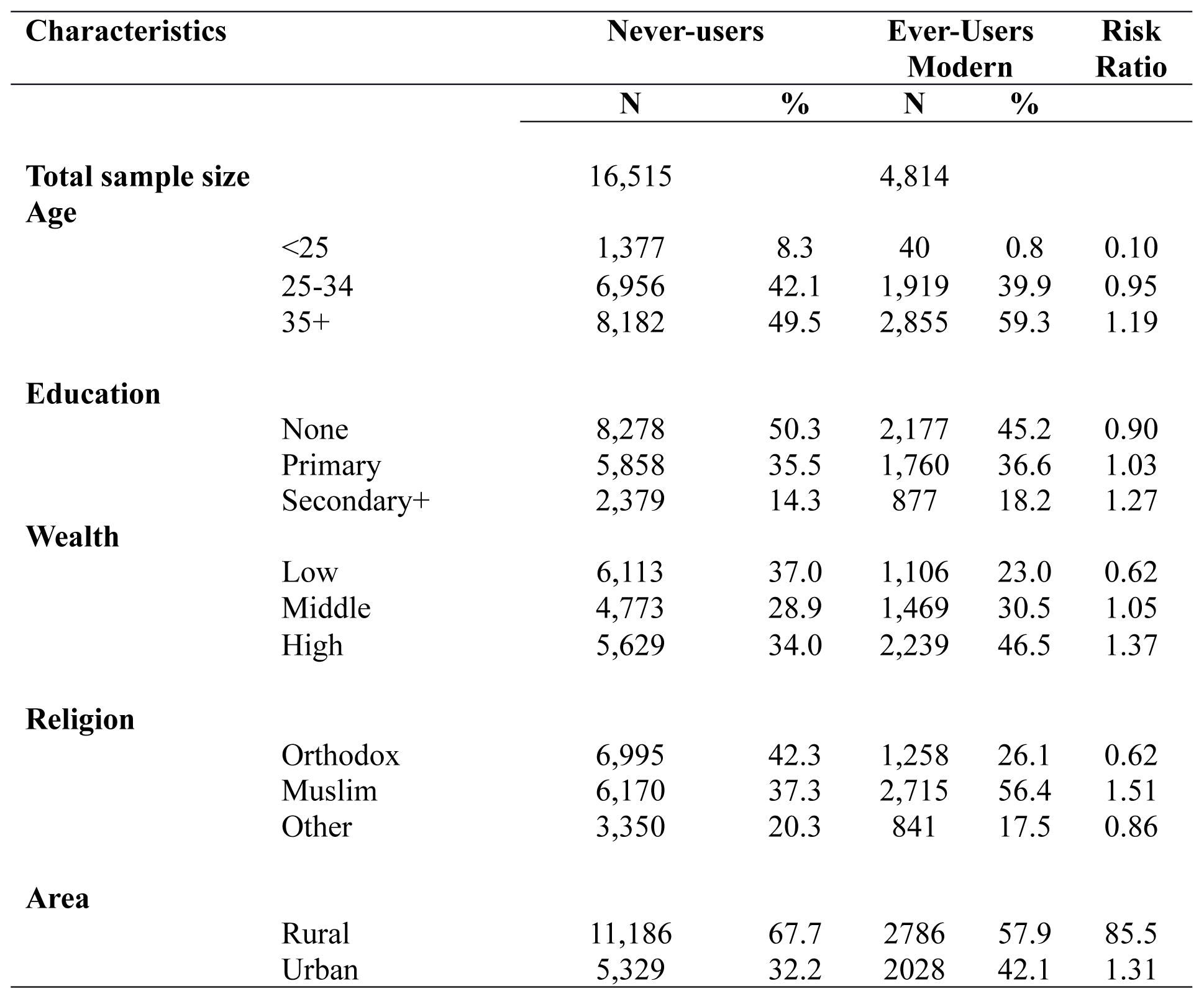
Description of the data. A risk ratio is computed to compare the subsample of women who have ever used contraception with the overall 2011 EDHS sample. Ever-users are on average older, more educated, wealthier, more urban and more likely to be Muslims compared to the overall EDHS data.

**Table 2:** Estimated standard Deviations and Pairwise Correlations for Woman-level Random Effects from the Multilevel models (2011)

| Random Effects | Discontinuation |  |  |
| --- | --- | --- | --- |
|  | Failure | Abandon | Switch |
| Failure | 1.14***<br>(0.18) |  |  |
| Abandon | 0.07<br>(0.25) | 1.08***<br>(0.08) |  |
| Switch | 0.52<br>(0.36) | 0.27<br>(0.24) | 1.02***<br>(0.18) |

The correlations between random effects are not significant therefore they are constrained to 0 in the following analyses. It means that hazards models can be modelled independently.

#### 3. Method choice

We used a multinomial probit model. Multinomial probit models are appropriate when the outcome of interest takes on a limited number of values that are not ordered. As compared to a logit model, it is particularly useful as it relaxes the hypothesis of independence of alternatives (IAA). Indeed, whether or not a woman will choose to use oral contraceptives might depend on the availability of condoms (another short-acting method). We thus ran a multinomial probit model with 4 possible alternatives (1) OC; (2) intra-uterine device (IUD) and implants; (3) injectables; (4) condoms. The way dependence between alternatives is considered is by correlating the residuals across the equations. Other methods of contraception are used in Ethiopia but we were only able to consider 4 choices due to the limitations of the aML software.

#### 4. Extension: multilevel multiprocess (allows for endogeneity)

Discontinuation and method choice were modelled jointly to allow for the fact that some women who are more likely to discontinue are also more likely to use short-acting contraceptives (i.e. the endogeneity of method choice). In effect, women level random effects are correlated between the discontinuation equations and the method choice equation [29]. Here we only considered 3 choices (OC, injectables and condoms, short-acting methods) due to software limitations.

### Interviews and focus group discussions (FGD)

Qualitative data were collected and analysed using a simple thematic approach to provide a deeper understanding of the major issues of enquiry generated from statistical results such as (1) Why do women discontinue contraceptive use in Ethiopia? (2) What sociocultural factors affect contraceptive discontinuation? (3) Why are short-acting hormonal methods both the most used and most discontinued? Thus the results from the interviews and FGDs have been summarised in a way to add further clarification to the statistical results.

Informed consent was obtained by all participants verbally before interviews and FGDs began. A written page to obtain informed consent was read out for each of the participants and the necessary explanation was given to each of the potential participants depending on the further clarification and additional information they required. Interviews and focus group discussions (FGDs) were conducted in 2013 in the three most populated regions in the country (the rural Arsi Administrative Zone of Oromia Region, the Sidama Administrative Zone of SNNP Region and the North Shewa Administrative Zone of Amhara Region). Qualitative data was collected on contraceptive discontinuation and method failure using 9 in-depth interviews and 6 FGDs in total among ever-married women of reproductive age (18-49) that had ever used contraception and 6 in-depth interviews among service providers (i.e. nurses and midwives) working in health facilities and health posts. These were distributed as 5 interviews per site (3 women and 2 service providers) and 2 FGDs with women per site. In all cases, only ever-married women were included as discussing sensitive issues related to sexual behaviour before marriage is not well accepted in such traditional societies of Ethiopia and only women over 18 were spoken to as that is the legal age of consent in Ethiopia.

In recruiting participants, inclusion criteria were selecting individuals of different socioeconomic class: poor, middle and better off individuals among residents in the community and of different levels of educational attainment: no-education, primary (1-8 grades) and secondary and above (grade 9 and higher achievers). Interviewees were approached at health facilities whilst visiting family planning (FP) clinics through the help of FP service providers or at their residence through the help of local persons serving as interview or FGD facilitators. There were no direct refusals to participate. However, some individuals were willing to take part in the study but were unable to, due to their busy schedule or inconvenience of time and location of the interview or FGD. The interviews then took place during an appointment at health facilities or at their residence depending on the preference of the interviewee. FGDs were done on school compounds or public places depending on availability and suitability of meeting places.

An experienced female research assistant conducted the interviews and FGDs, among women using local languages and lasting on average one and two hours, respectively. The interviews among service providers were conducted by the female research assistant or the last author, a male Ethiopian citizen working and living in the country, depending on location. The 2 service providers at each location consisted of 1 health extension worker (female) and 1 zone health office worker/supervisor (male or female) per location.

The questions asked in female interviews were as follows:

1. How often do you use contraception?
  a. What is the longest time you have used the same contraceptive continuously? Why?
2. Why do you use family planning methods?
  a. Could you tell me some of your specific reasons?
3. Where do you get the contraceptive supplies?
  a. Is it at your vicinity or at a distant location?
  b. Why do you prefer to go there?
4. *What type of contraception are you using?*
  a. *Could you tell me why you have chosen such a method?*
  b. *Was it the type of contraceptive you have used from the very beginning?*
5. *What challenges/problems were you facing in using contraception?*
  a. *Could you explain it to me further?*
  b. *How long have you faced such a problem?*
  c. *What measures have you taken to overcome such a problem?*

Focus group questions were similar but asked about the wider community’s contraceptive use and attitudes:

1. *How often do people in your community use contraception?*
2. *Why do people in your community use family planning methods?*
  a. *Could you tell us some of their specific reasons?*
3. *Where do people in your community get contraceptive supplies?*
  a. *Is it at their vicinity or at distant location?*
  b. *Why do you think they prefer to go to those places?*
4. *What type of contraception do people in your community often use?*
  a. *Could you tell us why they would choose such a method(s)?*
  b. *Is it the type of contraceptive they often take up from the very beginning?*
5. *What challenges/problems do women of reproductive age face in using contraceptives?*
  a. *Could you explain them further?*
  b. *What measures do they often take to overcome such a problem(s)?*

Service providers were asked the same questions as in the FGDs in their interviews with the addition of one extra question: What are the views of clients in using family planning methods? These interviews were conducted at respective workplaces of the service providers. All interviews and FGDs were tape recorded and transcribed in the language used for the interview and discussion. The summary of the major findings of the interviews and FGDs were then translated into English for inclusion in the paper.

## Results

### Trends of contraceptive discontinuation between 2000 and 2011

We compared the magnitude of contraceptive discontinuation between 2000 and 2011, using 5-years calendar data on contraceptive use from both the 2005 and 2011 Ethiopian DHSs. In the period 2006-11, the risk of discontinuation among women who use modern contraceptive methods is high: an episode of contraceptive uptake has a 56.4% risk of terminating, of which 64.6% correspond to abandonment, 15.9% correspond to failure and 19.5% correspond to switching. The overall level of discontinuation is slightly higher than in the period 2001-2005 (analysis of the 2005 EDHS not shown), and the persistence of a high level of discontinuation over the years might partly explain why the increase in contraceptive prevalence does not match the increase in the number of contraceptive ever-users (Figure 1).

**Figure 1.**
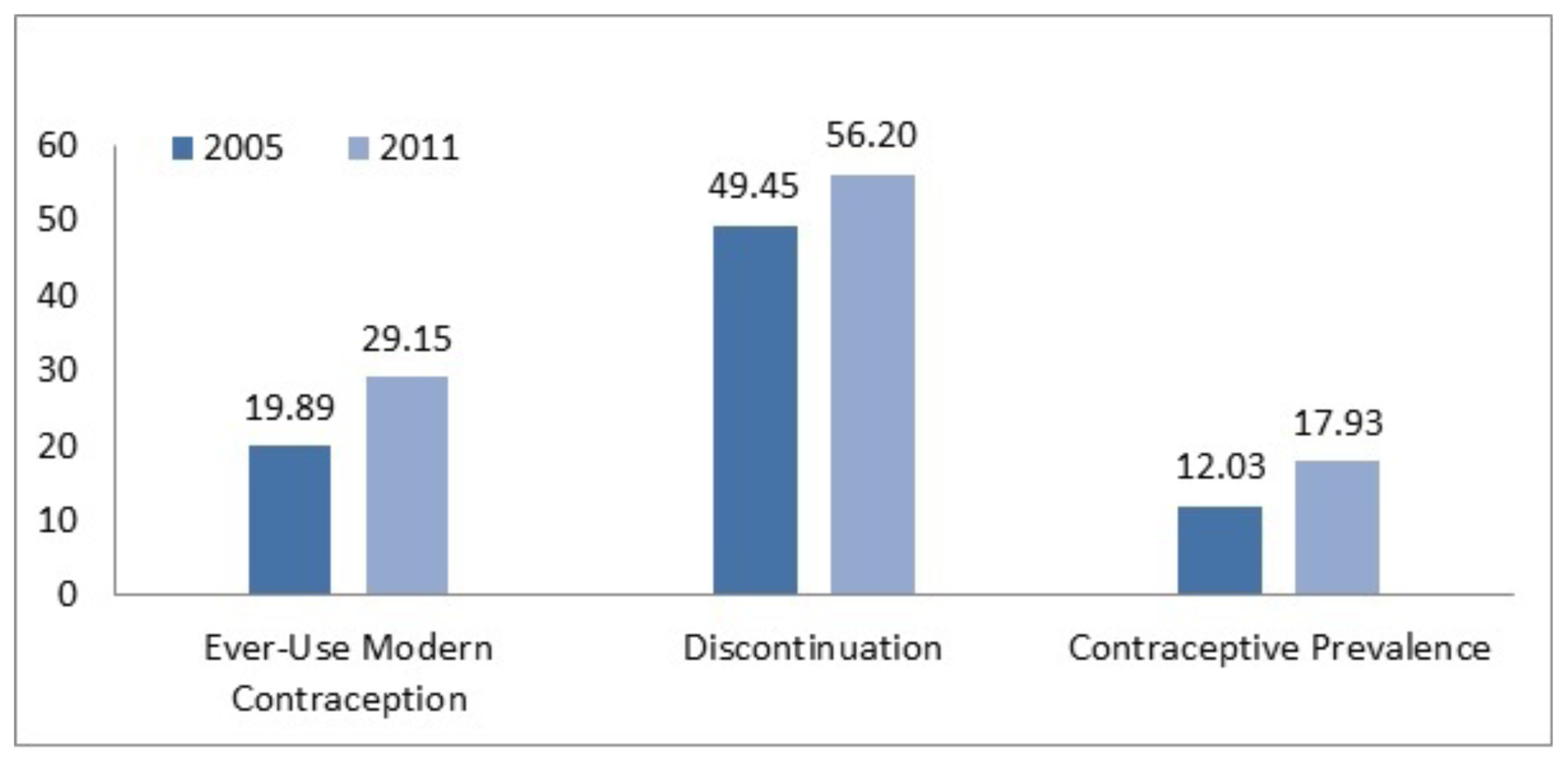
Trends in contraceptive adoption, discontinuation and use among Ethiopian women, 2005-2011. This figure is based on the descriptive statistics of the data collected during the 2005 and 2011 Ethiopian demographic and health surveys. The percentage of ever-users of modern contraception has increased by ca. 10% over the period investigated. The percentage of contraceptive discontinuation, which includes failure, switch and abandonment, has not reduced. The percentage of women using contraception at the time of the survey, i.e. contraceptive prevalence, has increased by ca. 5% in the study period.

### The determinants of contraceptive discontinuation

We investigated the hypothesis that contraceptive discontinuation is driven by the type of contraceptive method, but not education, among women who have ever-used contraception (see Table 2 for a description of the sample). First, we analysed method choice, and found that educated women are more likely to use oral contraceptives and condoms, while less educated women are more likely to use injectables and IUD/Implants (Table 3). The analysis controls for differences in age at the beginning of an episode of contraception, wealth, area, religion and patterns of correlation between methods types. Second, we analysed contraceptive discontinuation as a function of method type and socio-demographic variables, accounting for the effect of socio-demographic variables on the type of method used. We found that educated women are not more or less at risk of contraceptive abandonment and failure. However, educated women are more likely to switch methods (55% more likely than uneducated; OR=1.55; 95CI [1.05; 2.29]; Figure 2). Third, we found that the risk of all types of discontinuation is the highest for oral contraceptives (Figure 3): (1) Abandonment is most likely for the use of OC, compared to injectables (41% less likely than OC; OR=0.59; 95CI [0.49; 0.70]) and condoms (61% less likely than OC; OR=0.39; 95CI [0.26; 0.60]) (Table 4); (2) Failure is also more likely for the use of OC, compared to injectables (83% less likely than OC; OR=0.17; 95CI [0.13; 0.23]) and condoms (73% less likely than OC; OR=0.27; 95CI [0.13; 0.56]) (Table 5). (3) Switch, a form of discontinuation generally considered to be a marker of women’s ability to choose between methods and a phenomenon that can reduce unmet needs, is the most likely for oral contraceptives and the least likely for condoms (Table 6). Overall, contraceptive discontinuation is not driven by a lack of education but rather is a function of the type of method and other socio-demographic variables. All three types of discontinuation are more likely for the youngest (<25 years, Tables 4-6) and in the case of contraceptive abandonment, the risk is increased for both Muslims and the poorest women (Table 4).

**Figure 2.**
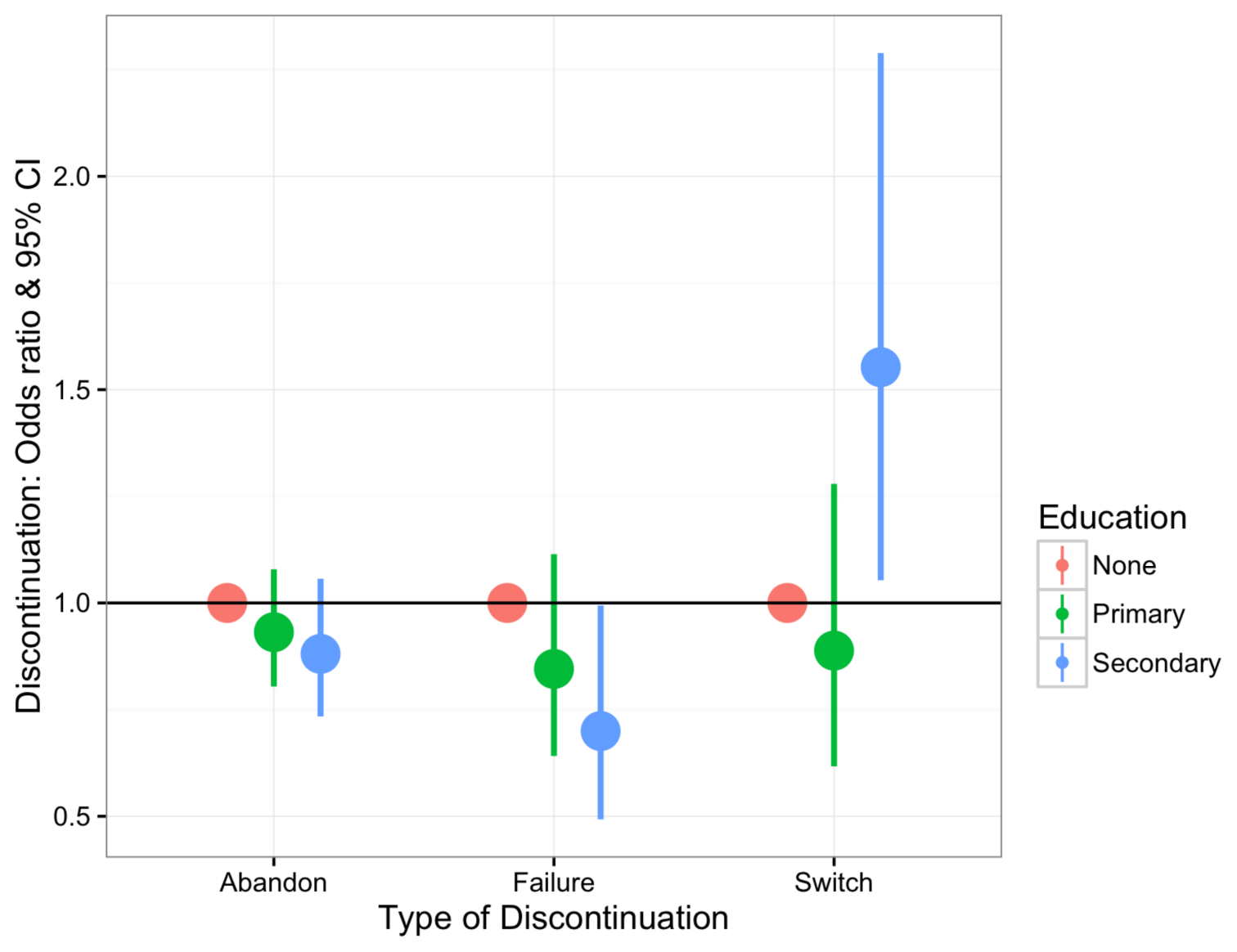
Predicted odd-ratios & 95% confidence intervals for contraceptive discontinuation as a function of a woman’s level of education. The reference category (horizontal line) is no education. The risk for a woman to discontinue contraception because of abandonment (a woman stops using contraception) or failure (a woman becomes pregnant) is independent from educational level. However, the risk of switching between methods is ca. 55% higher for women with the highest level of education. This figure is based on the results of a multilevel multi-process model using the calendar data collected during the 2011 Ethiopian demographic and health survey.

**Figure 3.**
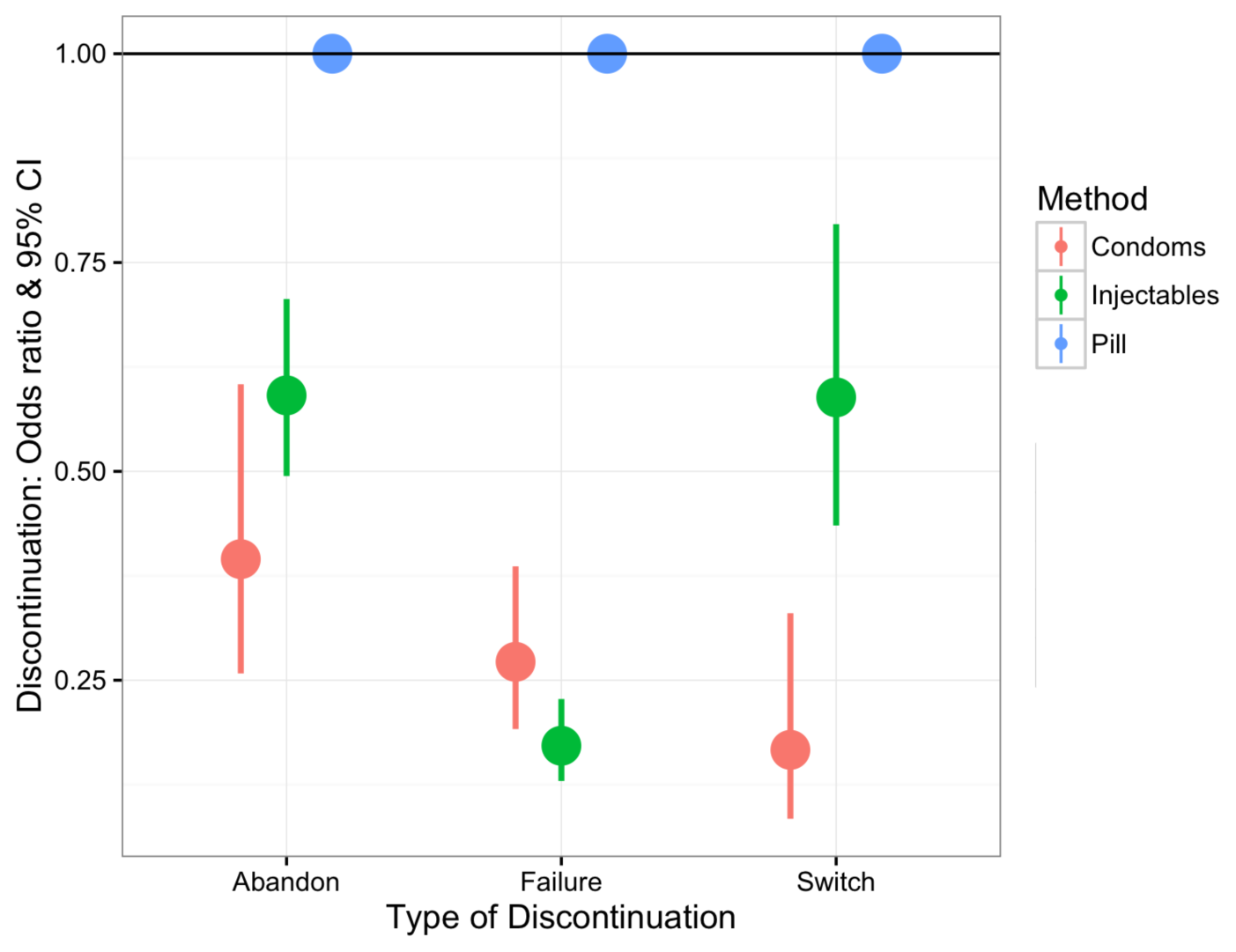
Predicted odd-ratios & 95% confidence intervals for contraceptive discontinuation as a function of method type. The analysis focuses on short-acting contraceptives only and the reference category (horizontal line) corresponds to the use of oral contraceptives. The risk of all three forms of discontinuation is higher for oral contraceptives. This figure is based on the results of a multilevel multi-process model using the calendar data collected during the 2011 Ethiopian demographic and health survey.

**Table 3:** Estimated Coefficients and Standard Errors for Multinomial Probit Model on Contraceptive Method Choice. The reference category is oral contraceptives. EDHS 2011.

| **METHOD CHOICE** | **<u>IUD/Implants</u>** |  | **<u>INJECTABLES</u>** |  | **<u>CONDOMS</u>** |  |
| --- | --- | --- | --- | --- | --- | --- |
| **Fixed Effects** | **Estimate** | **Std Err** | **Estimate** | **Std Err** | **Estimate** | **Std Err** |
| Constant | **-1.597\*\*\*** | **0.192** | **1.799\*\*\*** | **0.150** | **-2.176\*\*\*** | **0.400** |
| **Education** |  |  |  |  |  |  |
| No Education (Ref) | 0 | 0 | 0 | 0 | 0 | 0 |
| Primary | **-0.331\*\*\*** | **0.0777** | **-0.171\*\*\*** | **0.063** | 0.163 | 0.128 |
| Secondary | **-0.403\*\*\*** | **0.0844** | **-0.749\*\*\*** | **0.068** | 0.434 | 0.116 |
| **Age at start** |  |  |  |  |  |  |
| <25 years (Ref) | 0 | 0 | 0 | 0 | 0 | 0 |
| 25-34 years | **0.217\*** | **0.119** | -0.014 | 0.088 | **-0.460\*\*\*** | **0.141** |
| 35-49 | **0.560\*\*\*** | **0.119** | **-0.349\*\*\*** | **0.088** | **-0.891\*\*\*** | **0.147** |
| **Wealth** |  |  |  |  |  |  |
| Low | 0 | 0 | 0 | 0 | 0 | 0 |
| Medium | **0.394\*\*\*** | 0.139 | 0.149 | 0.120 | NA | NA |
| High | 0.026 | 0.142 | -0.097 | 0.117 | **0.897\*\*** | **0.395** |
| **Area** |  |  |  |  |  |  |
| Urban | 0 | 0 | 0 | 0 | 0 | 0 |
| Rural | **0.805\*\*\*** | **0.084** | **0.889\*\*\*** | **0.069** | **-0.447 \*\*\*** | **0.137** |
| **Religion** |  |  |  |  |  |  |
| Orthodox | 0 | 0 | 0 | 0 | 0 | 0 |
| Muslim | -0.006 | 0.070 | **-0.183\*\*\*** | **0.058** | **-1.232\*\*\*** | **0.131** |
| Others | 0.519 | 0.484 | 0.481 | 0.497 | NA | NA |
| **Correlation between women-level random effects across equations** |  |  |  |  |  |  |
| IUD/Im. & Injectables |  |  | **-0.097 (0.030) \*\*\*** |  |  |  |
| IUD/Im. & Condoms |  |  | -0.302 (0.188) |  |  |  |
| Condoms & Inject. |  |  | -0.077 (0.052) |  |  |  |

**Table 4:** Estimated Coefficients and Standard Errors for Hazards Models of Contraceptive Abandonment in 2011. The results for a standard model and multi-process model, modelling method choice conjointly, are compared. In the multi-process model, IUD is not considered due to the limitations of the aML software. Italics depict fixed parameters, estimated previously using a simple model with no variables.

| **ABANDON 2011** | **Standard model** |  | **Multi Process** |  |
| --- | --- | --- | --- | --- |
| **Variable** | **Coefficient** | **SE** | **Coefficient** | **SE** |
| **Constant** | *2.59* |  | -9.65 | 0.05 |
| **Duration (mths)** |  |  |  |  |
| 0-12 | *-0.723* |  | 0.226 |  |
| 12-24 | *0.560* |  | 0.129 |  |
| 24+ | *0.020* |  | 0.028 | 0 |
| **Method** |  |  |  |  |
| Pill | 0 |  | 0 |  |
| IUD/Implants | **-2.801\*\*\*** | **0.305** | -- | -- |
| Injectables | **-1.512\*\*\*** | **0.270** | **-0.526\*\*\*** | **0.091** |
| Condoms | -0.069 | 0.144 | **-0.929\*\*\*** | **0.217** |
| **Education** |  |  |  |  |
| No Ed | 0 |  | 0 |  |
| Primary | -0.559 | 0.337 | -0.071 | 0.075 |
| Secondary | 0.079 | 0.121 | -0.127 | 0.093 |
| **Age at start** |  |  |  |  |
| <25 | 0 |  | 0 |  |
| 25-34 | 0.044 | 0.149 | **-0.460\*\*\*** | **0.100** |
| 35-49 | -0.148 | 0.1716 | **-0.947\*\*\*** | **0.101** |
| **Wealth** |  |  |  |  |
| Low (1-2) | 0 |  | 0 |  |
| Medium (3-4) | **-0.498\*\*** | **0.1814** | **-0.360\*\*\*** | **0.111** |
| High (5-6) | -0.186 | 0.1975 | **-0.539\*\*\*** | **0.102** |
| **Area** |  |  |  |  |
| Urban | 0 |  | 0 |  |
| Rural | -0.334 | 0.1982 | 0.0007 | 0.0812 |
| **Religion** |  |  |  |  |
| Orthodox | 0 |  | 0 |  |
| Muslims | 0.1658 | 0.136 | **0.175\*\*** | **0.076** |
| Others | 0.226 | 0.1156 | -0.225 | 0.442 |
| **Women level effects** |  |  |  |  |
| Sigma | 0.30 | 0.24 | **0.406\*\*\*** | **0.1063** |
| Rho (abandon/injectable) | -- | -- | **0.2149\*\*** | **0.0935** |
| Rho (abandon/condom) | -- | -- | 0.2290 | 0.1486 |
| Rho (injectable/condom) | -- | -- | **-0.0917\*\*\*** | **0.0353** |

**Table 5:** Estimated Coefficients and Standard Errors for Hazards Models of Contraceptive Failure in 2011. The results for a standard model and multi-process model, modelling method choice conjointly, are compared. In the multi-process model, IUD/Implants are not considered due to the limitations of the aML software. Italics depict fixed parameters, estimated previously using a simple model with no variables.

| **FAILURE 2011** | **Standard model** |  | **Multi Process** |  |
| --- | --- | --- | --- | --- |
| **Variable** | **Coefficient** | **SE** | **Coefficient** | **SE** |
| **Constant** | *9.82* |  | *-9.92* | *0.18* |
| **Dur12** | *-0.1557* |  | *0.222* |  |
| **Dur24** | *0.228* |  | *0.123* |  |
| **Dur36** | *-0.009* |  | *-0.001* |  |
| **Method** |  |  |  |  |
| Pill | 0 |  | 0 |  |
| IUD/Implants | **1.834\*\*\*** | **0.365** | -- | -- |
| Injectables | **-2.9592\*\*\*** | **0.5317** | **-1.763\*\*\*** | **0.144** |
| Condoms | **-1.5319\*\*\*** | **0.1518** | **-1.302\*\*\*** | **0.369** |
| **Education** |  |  |  |  |
| No Education | 0 |  | 0 |  |
| Primary | **-0.9710\*\*** | **0.3471** | **-0.168** | **0.141** |
| Second | -0.111 | 0.1458 | -0.357 | 0.179 |
| **Age Start** |  |  |  |  |
| <25 yrs | 0 |  | 0 |  |
| 25-34 yrs | -0.2544 |  |  |  |
|  |  | 0.1875 |  |  |
| 35-49 yrs | **-0.5526\*\*** | **0.2046** | **-1.42\*\*\*** | **0.188** |
| **Wealth** |  |  |  |  |
| Low | 0 |  | 0 |  |
| Medium | **-1.326\*\*\*** | **0.2161** | 0.169 | 0.219 |
| High | 0.258 | 0.2607 | -0.301 | 0.200 |
| **Area** |  |  |  |  |
| Rural | 0 |  | 0 |  |
| Urban | -0.160 | 0.266 | 0.201 | 0.164 |
| **Religion** |  |  |  |  |
| Orthodox | 0 |  | 0 |  |
| Muslim | 0.2770 | 0.1769 | 0.409 | 0.136 |
| Other | **0.4770\*\*\*** | **0.1352** |  |  |
| **Women level effect** |  |  |  |  |
| Sigma | 0.43 | 0.62 | 0.489 | 0.133 |
| Rho (failure/injectable) |  |  | 0.142 | 0.110 |
| Rho (failure/condom) |  |  | 0.170 | 0.155 |
| Rho (injectable/condom) |  |  | **-0.097\*\*\*** | **0.035** |

**Table 6:** Estimated Coefficients and Standard Errors for Hazards Models of Contraceptive Switch in 2011. The results for a standard model and multi-process model, modelling method choice conjointly, are compared. In the multi-process model, IUD/Implants are not considered due to the limitations of the aML software. Italics depict fixed parameters, estimated previously using a simple model with no variables.

| **SWITCH2011**<br>**Variable** | **Standard Model** |  | **Multi Process** |  |
| --- | --- | --- | --- | --- |
|  | **Coefficient** | **SE** | **Coefficient** | **SE** |
| **Constant** | *9.81* |  | -9.92 | 0.18 |
| **Duration (mth)** |  |  |  |  |
| 0-12 | *-0.189* |  | 0.19 |  |
| 12-24 | *0.255* |  | 0.15 |  |
| 24+ | *0.011* |  | 0.01 |  |
| **Method** |  |  |  |  |
| Pill | 0 |  | 0 |  |
| IUD/Implants | 0.399 | 0.412 | -- | -- |
| Injectables | -0.436 | 0.249 | **-0.530\*\*\*** | **0.154** |
| Condoms | **-0.646\*\*\*** | **0.155** | **-1.792\*\*\*** | **0.349** |
| **Education** |  |  |  |  |
| No Ed | 0 |  | 0 |  |
| Primary | -0.67 | 0.346 | -0.1184 | 0.186 |
| Secondary | -0.072 | 0.166 | **0.440\*\*\*** | **0.198** |
| **Age at start** |  |  |  |  |
| <25 | 0 |  | 0 |  |
| 25-34 | 0.356 | 0.178 | -0.2599 | 0.2303 |
| 35-49 | -0.379 | 0.243 | **-0.517\*\*\*** | **0.235** |
| **Wealth** |  |  |  |  |
| Low (1-2) | 0 |  | 0 |  |
| Medium (3-4) | **-0.6417\*\*** | **0.2510** | -0.041 | 0.280 |
| High (5-6) | -0.016 | 0.311 | 0.336 | 0.229 |
| **Area** |  |  |  |  |
| Urban | 0 |  | 0 |  |
| Rural | .0356 | 0.305 | 0.156 | 0.177 |
| **Religion** |  |  |  |  |
| Orthodox | 0 |  | 0 |  |
| Muslim | 0.235 | 0.1624 | 0.029 | 0.1699 |
| Other | 0.047 | 0.152 | -- | -- |
| **Women level effects** |  |  |  |  |
| Sigma | **1.029\*\*** | **0.39** | **0.489\*\*\*** | **0.137** |
| Rho (failure/injectable) |  |  | 0.142 | 0.110 |
| Rho (failure/condom) |  |  | 0.170 | 0.155 |
| Rho<br>(injectable/condom) |  |  | **-0.097\*\*\*** | **0.035** |

### Barriers to continuation

#### 1. Side-effects: *“As I started to take the injectable, I ended up with much bleeding”*

High levels of abandonment may be due to method-related reasons such as physiological side effects and health concerns, associated especially with hormonal methods. Focus group discussion participants commonly cited that the major reason for discontinuation of hormonal contraceptives is the ‘excessive bleeding that women encounter at the beginning or after sometime of using the methods’. One mother of 3 from Arsi village with primary level education stated that “As *I started to take the injectable, I ended up with much bleeding [menstrual irregularities]*.” Bleeding is considered as a major side effect of contraceptive use that could lead to loss of life as a result of getting weaker due to losing too much blood from the body. In addition, the literal meaning of the term used to describe ‘excessive bleeding’, ‘ dem bizat’, can be translated back into ‘too much blood’ in Amharic, is wrongly connoted with the effects of high blood pressure that can lead to an immediate death.

Other mentioned side effects as well as bleeding that women experienced that led them to discontinue hormonal methods included pain, dizziness, skin conditions and mood and appetite changes. For instance, although not included in the quantitative analysis, another hormonal method, the implant (commonly referred to in Ethiopia as ‘bars’) was cited by several women as being discontinued due to health concerns. For example, a married woman living in a small town in southern Ethiopian, with secondary education and two children, said that in the first few days after receiving the implant she felt restless and dizzy with severe and constant pain in her left arm and she was unable to hold things up in that hand. Having suffered from pain and psychological tension for nearly three months, she decided to have it removed. When asked about her main worry that led her to have it removed, she said that: “… *I do not want to lose my muscles. For me, it is better to have as many children as God let me, than getting paralyzed having gone against God’s will*”. Side effects reported by other women include “*Pills bring about irritation [scars] on the face*” (uneducated, 39, mother of 5, Arsi village) and “*the loop [IUD] brings about excessive bleeding and loss of appetite*” (primary school, 27, mother of 3, North Shewa small town).

Many participants, if they had not experienced side effects themselves, had heard rumours and reports of other women’s side effects which may make them more likely to abandon contraception themselves. Health concerns and rumours expressed by women in our study include: “*Taking pills causes stomach-ache due to the chemical reaction to our body cells*” (primary school achiever, 37, Sidama small town, mother of 3); “*Taking pills burns a woman’s face due to its chemical reaction in the body*” (uneducated women, 42, mother of 6, Arsi village); *“Pills can be concentrated in the womb and damage the woman’s reproductive organ*” (primary school achiever, 24, mother of 3, North Shewa village), “A *woman should not use injectables until she achieves her ideal family size as it cause secondary sterility*” (uneducated, 38, mother of 5, Arsi rural village.) Although not in all cases, many of these rumours appear to have a basis in the reports of what women believe they have actually experienced that then sometimes become amplified, distorted and wrongly explained through several occurrences of social transmission thus many ‘misconceptions’ may have some grounding in real experience.

#### 2. Poverty

We investigated the possibility that women living in poverty with worse living conditions experience more severe side-effects due to being less able to tolerate exogenous hormones doses. This was supported by both the quantitative and qualitative findings. The qualitative approach revealed that many women believe that hormonal contraception is less tolerable for women living in poverty and that these women are more likely to abandon hormonal contraception. The quantitative analysis showed that abandonment, but not failure or switch, is associated with poverty: it is 42% less likely among the wealthiest (OR=0.58; 95CI [0.48, 0.71], Table 4). Indeed, we found that wealth was the most important predictor, along with method type, for predicting the odds of contraceptive abandonment (Figure 3). There was a perception among women in the qualitative interviews that hormonal contraception is not well suited to the bodies of women in lower socioeconomic classes with a poor diet. For instance, one woman (primary school achiever, 32, mother of 3, North Shewa small town) stated that “*Only women having access to better diet [egg, meat, butter…] should take pills or injectables as it does not work for those having poor diet.… the tablets [pills]/ dose [injectables] need better diets to operate in the woman’s body*” and others report fears of hormonal contraceptives using up energy from their already small food intake and making them hungry or ill. Participants who underwent excessive bleeding reported abandoning contraception because they were told that their body was getting weaker and that they needed to take a balanced diet to replace the blood that they had already lost. Specifically, women living in poverty often took the immediate action of stopping the method as the suggested solution was beyond their economic reach, and was their greatest concern. These findings suggest that side effects may be experienced to a greater extent or felt more severely in the lower socioeconomic statuses, leading to higher levels of abandonment.

#### 3. The need for secrecy

A barrier to continuation commonly cited in the interviews is the fear of a woman’s contraceptive use being found out by her husband. Many women reported that their husbands disapproved of contraception and associated it with infidelity. Therefore, if a woman experiences physiological side effects, it can compromise the secrecy of her contraceptive use which might force her to discontinue. The need to keep contraceptive use a secret also hinders a woman’s ability to seek help or advice for side effects by contacting the family planning service providers. Indeed, women reported that to visit the nearby health facility to speak to a health care provider or get their resupply, they needed to have a convincing reason to avoid the suspicion of their husbands. As this was often not possible, women were more likely to abandon contraception. The account from one married rural woman with primary level education and mother of 4 children sums up well what women cited as to why their husbands were suspicious of the use of contraception.

“*I started using pills without the knowledge of my husband as he was not in favour of family planning services for his mental setting is to associate contraceptive use with sexual infidelity. He thinks that women are using contraceptives to enhance their chances of ‘cheating on their husbands’ as it prevents pregnancy that could occur outside marriage*.”

Raised suspicion from the husband for any reason and fear of him finding out, may cause a woman to have to abandon her use of contraception. It seems for many women there is no solution which does not completely eliminate this risk, as shown by the experiences of the same married rural woman with primary level education and mother of four children:

“*I decided to take oral pills to prevent pregnancy. As I had been taking the pills every night before going to bed (actually hiding it outside a bedroom) and drinking water to swallow it, he [husband] got suspicious of my act and began to question why I do that every night regularly. Hence, I decided to use another method that would not get him clue of my act to prevent pregnancy. Then I decided to switch to another method, the injectables initially. However, I found it difficult to do so as it requires visiting the nearby health post or health center at least every three months for which I could not come-up with ‘good reasons’ to do so regularly. My intention to use implants did not also work out as the scars on my arms could alert the suspicion of my husband. Whilst consulting my closest friend living in a nearby town and seeking advice from her, I learnt that an IUD is the safest as it can stay for years without necessarily going through resupply every time. Though I have successfully got the IUD inserted, I got to be worried about my husband’s complaints of “sexual pleasure” since recently. He repeatedly complained that he feels something strange in my body during sexual consummation, and asked if I did anything that is unknown to him. Having realized that his question was emanating from the discussion he had with his male friends that do not support family planning services, I decided to remove the IUD to save my marriage and maintain* ‘*peace*’ *in my marital life and the wellbeing of our children*.”

Even among women themselves, there is some sense of association between use of contraception and infidelity with opinions being expressed such as: “*Women who are loyal to their husbands should not use condom*” (primary school, 26, mother of 1, North Shewa village) and “*The idea of using condom creates suspicion between the wife and the husband*” (secondary school, 33, mother of 5, Arsi small town.) However even if a woman does not hold these associations and wishes to continue taking contraception to limit or space her births, it may be very hard for her to do so if she fears the consequences of her husband finding out she is taking it. This in part explains the popularity of injectables as it affords secrecy by leaving no trace on the body and only needs to be administered once every 3 months. Nationally, women are 6 times more likely to use injectables compared to oral contraceptives (OR: 5.99; 95CI [4.48; 8.08]) (Figure 4). Above all, the educational campaign held in the country in the late 1990s to make injectables popular made a significant contribution to the wider use of the methods.

**Figure 4.**
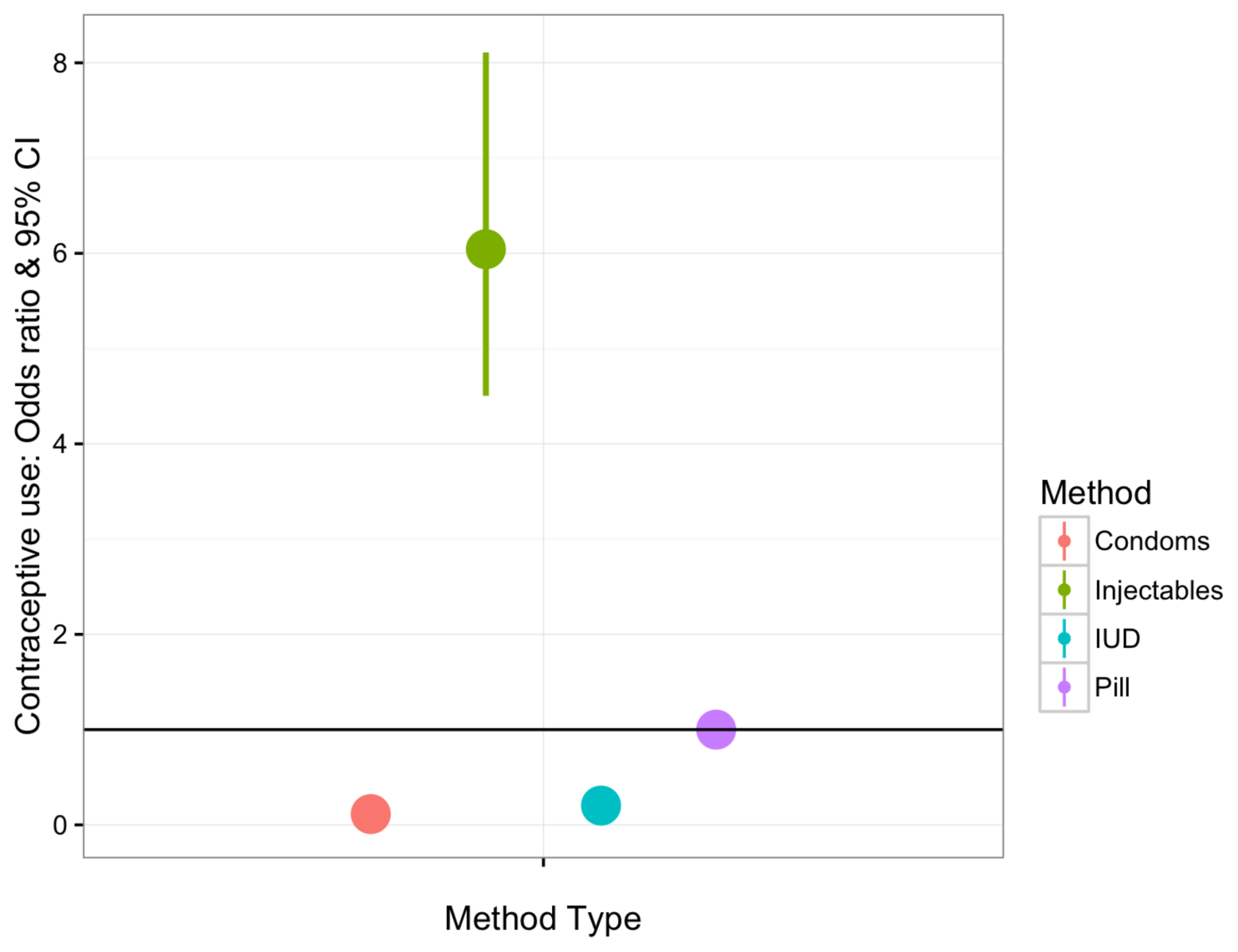
Predicted odd-ratios & 95 CI for method choice. The reference category (horizontal line) corresponds to the use of oral contraceptives. At the national level, injectables are 6 times more likely be used than any other short-acting methods. This figure is based on the results of a multinomial probit model using the calendar data collected during the 2011 Ethiopian demographic and health survey.

## Discussion

Contraceptive discontinuation has been identified as a major challenge to tackling unmet needs of contraception and unwanted fertility in the developing world [7–9]. This paper aims to provide a better understanding of the reasons why women discontinue contraception in Ethiopia, a country where 1 in 5 women faces unmet need for contraception (DHS data) despite a nine-fold increase in the number of women who have ever used contraception. Combining an analysis of the 2011 EDHS data with semi-structured interviews with Ethiopian women and health professionals in 2013, we found that (1) overall 36 months rates of discontinuation have not decreased in the period 2001-2011 (from 49.4% in the 2005 EDHS (analysis not shown) to 56.4% in the 2011 EDHS), (2) discontinuation is significant and multifactorial, and mainly takes the form of contraceptive abandonment (2/3 of all discontinuation rates) as compared to switch and failure and (3) the main barriers to continuation are (i) the experience of physiological side-effects associated with the use of hormonal contraceptives, (ii) the conflation of contraceptive use with the occurrence of marital infidelity, and (iii) poverty. Taken together, our results show that female education is not a barrier to contraceptive continuation, and suggest a shift in perspective focusing on increasing male education and questioning the appropriateness of medical technology to the physiology of Ethiopian women, especially those living in poverty.

### Risk of discontinuation, method type and wealth

The overall 36 months risk of discontinuation of modern contraception is just over 50%, and out of the three forms of contraceptive discontinuation studied, i.e. abandonment, failure, and method switch, abandonment accounts for 2/3 of all discontinuation rates. Contraceptive abandonment does not always lead to unwanted pregnancy, however, as there are many reasons why women might stop using contraceptives, e.g. desire for more children, no further needs, method-related reasons and health concerns/side-effects. In the 2005 Ethiopia DHS, Ali, Cleland and Shah found that no further need and the desire for another child accounted for 15% and 33%, respectively, of the overall rate of contraceptive discontinuation [8]. This suggests that although abandonment is not always the results of method-related reasons and health concerns, those reasons account for a significant part of the overall rate of discontinuation in Ethiopia. The 2011 data does not allow quantifying the contribution of each type of reason to the overall abandonment rate, but our analysis shows that the risk of abandonment is a function of the type of method used. Of particular interest is the finding that oral contraceptives are ca. 40% more likely to be abandoned than injectables and condoms. This picture differs from the results obtained in a previous analysis of 19 developing countries, where the 12 months probability to discontinue due to method-related reasons, side-effects or health concerns is found to be the highest for injections compared to the pill [8]. A possibility for explaining the discrepancy between the two analyses is that the statistical method used in this paper (i.e. a multiprocess multilevel model [29]) accounts for the effect of socio-demographic variables on the type of method used when quantifying the impact of the type of method used on the risk of abandonment. Indeed, users of injectables are more likely to be the poorest and the poorest are those who face the highest risk of contraceptive abandonment. Why poverty leads to an increased risk of abandonment may be due to a stronger desire for more children among the poorest, and/or the experience of more side-effects. An additional possibility is that injections and oral contraceptives are both associated with side-effects, but that injectables afford a benefit oral contraceptives do not (e.g. the ability of the method to afford secrecy).

### Contraceptive use and the need for secrecy

One determinant of discontinuation that emerged as critical in reducing women’s ability to continue taking contraception, switch method or seek advice, was the association of contraceptive use with infidelity and thus the need for keeping contraceptive use secret. Fear of husband’s suspicion of contraceptive use might go some way to explaining the popularity of injectables as it leaves no trace on the body and, compared to OC, it only needs to be administered once every 3 months thus minimizing the number of visits to the health centre. The difference in how much secrecy OC and injectables afford may explain why the injectables are less abandoned and switched than OC. Although injectables are the most commonly used method - a quarter of married women and a third of unmarried sexually active women use injectables (EDHS 2016) – there remains unnecessarily high levels of abandonment due to reported side effects that both compromise the secrecy of its use and are understood as a serious health threat.

### The experience of physiological side-effects

The qualitative data indicate that women experience debilitating side-effects in the form of excessive bleeding, which is interpreted as a threat to a woman’s health and her ability to conceive. Given a woman’s capacity to reproduce has traditionally been a critical asset in her marital union [31], the experience of side-effects further puts women at risk with their marriage. The findings are in line with a recent report from the UNFPA suggesting that the main reasons cited for not using “family planning worldwide are the fear of side effects and health concerns” [32]. While the literature on side-effects is generally unclear with regards to the distinction between “real” side-effects and unsubstantiated rumours [33, 34], our findings indicate that the fear of many types of side-effects is likely to originate in the experience of users. Why women experience physiological side-effects might result from a mismatch [13] between the dose of hormones found in the contraceptives used in Ethiopia and the physiology of Ethiopian women. Although a clear general pattern is difficult to draw, especially when taking within populations comparisons [20], levels of ovarian steroids are often found to be higher for women living in rich environments and lower for women living in ecologies characterized by high physical workload [35] and nutritional stress [20, 36]. For instance, as compared with women living in the US, salivary progesterone are significantly lower for women living in Zaire [37], Nepal [38] and Poland [35]. Thus, our findings that abandonment and the experience of side effects are more likely to occur for women who are living in poverty are in line with the hypothesis according to which side-effects result from a mismatch between local hormonal levels and those present in the contraceptives obtained from international donors. If further analysis reveals that to be the case, then a focus on tackling the cause of side-effects through developing new contraceptives adapted to the local reproductive ecology may have drastic consequences for decreasing unmet needs for contraception.

### Education and contraceptive discontinuation

The results show that less educated women are not more likely to abandon or fail using modern contraception, suggesting that educational programs, on their own, might not be able to tackle the roots of discontinuation. Rather, we found that better educated women were more likely to switch methods, which is in line with previous reports [39, 40]. For instance, a review of 6 developing countries found that pill continuation was overall not correlated with a woman’s level of education and concluded that counselling for uneducated women was not in need of radical review [41]. It was also previously found that switch, but not failure or abandonment, was linked to education, with probability of switching rising with levels of education [39]. Our results support these findings and contrast with calls for more information through education and counselling to reduce discontinuation, especially abandonment. For instance, to reduce high rates of discontinuation, it is recommended that women should be “forewarned about side-effects and reassured about health concerns” [8] and another report suggests “dispelling misconceptions and counselling women who experience amenorrhea” [9]. While information is undoubtedly mandatory, it is not sufficient. In a country such as Ethiopia where contraception is frequently used for spacing births [42, 43], it is no surprise that the experience of physiological side-effects that weaken the body is a major barrier to continuation. It has been shown that in sub-Saharan Africa, contraceptives may be used to heal and maintain strength in the face of repeated pregnancies and reproductive mishaps, with the decline in bodily resources being seen as the most significant barrier to achieving high fertility [44]. It follows that the recommended best practice in the face of side-effects - “If a client experiences common side-effects […], you should advise the woman to keep taking her pills” [45] – may be beyond the reach of most women as their experience is too debilitating [19].

### Implications

The results do confirm the importance of men’s opinion of family planning and the need for informative, confidential service provision. A report on reducing discontinuation suggests that engaging male partners by “enhancing couple communication about method characteristics can be effective in supporting continued use, particularly in the postpartum period” [9]. Interventions targeting male leaders, husbands and men might indeed allow women to seek counselling about any side effects and get resupplies closer to home. However, this approach focuses on information and social cognition in determining behaviour, missing out the roles of both past behaviour or habituation and non-cognitive determinants such as social norms [46]. Social norms around marriage and reproduction might be resistant to educational programs in the absence of ecological change shaping the demand for children (e.g. variation in the opportunity cost of children due to lower mortality rates and/or stronger links between mother’s education and wealth [47]. Therefore, in the meantime, improving the experience of methods that afford women the secrecy they need may be more effective at reducing discontinuation.

As secrecy is still required, injectables provide the best current solution for keeping a woman’s contraceptive use secret. Yet, injectables are still commonly abandoned by women who become at risk of unwanted pregnancy due to the experience of side effects that can both jeopardise the maintenance of secrecy and be too debilitating to enable the continuation of contraceptive use. Thus, pharmaceutical companies should be engaged to develop contraceptives more suited to the physiology of women living in different ecologies that will lessen side effects. A new lower-dose self-administered injectable, the Sayana Press, is now licensed in many countries and may prove more tolerable due to its lower dose. However, this product was tested in U.S. and Singaporean women, in samples with a mean BMI of 28.7 and 22.4 respectively [48], and comes with special warnings about high risk of menstrual irregularities and loss of bone mineral density. Further, the US sample was compromised of 41% obese women (with BMIs >30) who are likely to have physiologies far removed from that of a rural Ethiopian woman living in poverty. Thus there is a need of trialling contraceptives on populations of women living in different ecologies before determining their acceptability.

The Ethiopian Federal Ministry of Health focus their policies on dispelling ‘myths’. For instance, the online resource available for family planning counselling with regards to oral contraceptive pills [45] begins with listing common “myths”: “Women who stop taking the pill may not be able to get pregnant; They become infertile; The pill causes cancer; Oral pills build up in a woman’s body; Women need to rest from taking oral contraceptives on sex-free days; Oral contraceptives cause birth defects or multiple births; Oral contraceptives change women’s sexual behaviour; Oral contraceptives accumulate in a woman’s stomach”. While misconception is perceived as the most pressing issue, it is only a few pages later that the most common side-effects, i.e. changes in menstrual bleeding patterns (irregular periods, spotting, amenorrhoea), headache, nausea, sore breasts, mood changes and sometimes high blood pressure, are covered. Our research suggests a shift away from conceptualising side effects as rumours or minor problems to acknowledge real problematic and sometimes intolerable experiences. Service providers would benefit from being educated to distinguish between substantiated concerns (i.e. irregular bleeding) and myths when evaluating whether to tackle misconception and when to counsel switch to a different method and/or other solutions. Note however that many of the interventions that may help alleviate some of the worst side effects, such as painkillers or iron supplements to replace that lost through irregular bleeding, are out of the reach of many women in Ethiopia.

### Limitations

We have used both semi-structured interviews and DHS data but the information collected in both cases is not directly comparable. Indeed, the quantitative data have been collected 2-years before the interviews were conducted, and the sample of women interviewed likely represents a sub-set of the nationally representative DHS data. For instance, only ever-married women were included in the qualitative analysis due to the cultural unacceptability of discussing sensitive issues related to sexual behaviour before marriage. It follows that not all reasons for discontinuation have possibly been voiced. This combined with the absence of data on the reasons for discontinuation in the 2011 EDHS data means that the quantification of the contribution of each of the possible reasons to discontinuation rates is beyond the scope of this paper. Yet, the findings that (i) at the national level, discontinuation depends on the type of contraceptive method and (ii) among married women in need of contraception, the experience of physiological and social side effects promotes contraceptive discontinuation, together point towards method type as a root cause for discontinuation. In the absence of physiological data, the use of mixed methods has helped direct attention towards to possible role of a mismatch between a woman’s endogenous hormonal levels and the exogenous dose of hormones given in contraception. Further research should explore whether Ethiopian women with lower levels of endogenous hormones are disproportionately affected by side effects.

## Conclusions

In line with recent reports on improving family planning programs worldwide [9, 32], we conclude that for Ethiopia, a strategy prioritizing the tackling of discontinuation due to side-effects over the increasing of the number of new clients may have drastic consequences for decreasing unmet needs for family planning. Further, to tackle discontinuation due to side-effects, we argue that dispelling misconceptions through educating women is not addressing the root causes of discontinuation, and that priority should be given to both engaging men and tackling side-effects by developing new contraceptives that are physiologically better adapted to the population.

## List of abbreviations

BMI: Body Mass Index
95CI: 95% Confidence Interval
DHS: Demographic and Health Survey
EDHS: Ethiopian Demographic and Health Survey
FGD: Focus Group Discussion
FMOH: Federal Ministry of Health
FP: Family Planning
IAA: Independence of Irrelevant Alternatives
IUD: Intra Uterine Device
OC: Oral Contraceptives
OR: Odd-Ratio
SNNP: Southern Nations, Nationalities, and Peoples′ Region
TFR: Total Fertility Rate
UNFPA: United Nations Population Fund (United Nations Fund for Population Activities)
U.S.: United States

## Declarations

### Ethics approval and consent to participate

For the qualitative study, ethic approval has been granted by the Institute of Development and Policy Research from the Ethiopian University of Addis Ababa. The ethical procedure associated with DHS data collection can be found on the DHS website http://dhsprogram.com/What-We-Do/Protecting-the-Privacy-of-DHS-Survey-Respondents.cfm

### Consent for publication

Oral consent was obtained for publishing anonymized quotes, as part of the general ethical procedure.

### Availability of data and material

DHS data can be retrieved for free from the DHS website http://dhsprogram.com. Qualitative data are available from the corresponding author on reasonable request.

### Authors′ contributions

EG designed the study, EG and AA collected qualitative data, EG analyzed the qualitative data, AA analyzed the quantitative data, AA, RS and EG wrote the paper.

## Acknowledgements

We are very grateful to all participants who answered our questions as part of our qualitative investigation.

